# Comparing Inference under the MSC versus the MSC-with-Recombination

**DOI:** 10.1101/2022.09.03.506495

**Authors:** Zhi Yan, Huw A. Ogilvie, Luay Nakhleh

## Abstract

Accurate inference of population parameters plays a pivotal role in unravelling the evolutionary histories. While recombination has been universally accepted as a fundamental process in the evolution of sexually reproducing organisms, it remains challenging to model it exactly. Thus, existing coalescent-based approaches make different assumptions or approximations to facilitate phylogenetic inference, which can potentially bring about biases in estimates of evolutionary parameters when recombination is present. In this article, we evaluate the performance of population parameters estimation using three classes of methods: StarBEAST2, SNAPP, and diCal2. We performed whole-genome simulations in which recombination rates, mutation rates, and levels of ILS were varied. We show that StarBEAST2 using short or medium-sized loci is robust to realistic rates of recombination, which is in agreement with previous studies. SNAPP, as expected, is generally unaffected by recombination events. Most surprisingly, diCal2, a method that is designed to explicitly account for recombination, performs considerably worse than other methods under comparison.

## 1 Introduction

The development of statistical models has moved phylogenetics beyond the qualitative estimates of cladistic relationships to inference of demographic histories including divergence times, population size changes and other quantitative values. These statistical methods rely on the fact that different loci across the genome have distinct histories which arise from recombination during sexual reproduction.

Topological discordance between gene and species phylogenies resulting from this recombination is known as incomplete lineage sorting (ILS). The multispecies coalescent (MSC) model accounts for this discordance (Hudson, 1983; Rannala and Yang, 2003), and because ILS is often the primary source of gene tree heterogeneity, the MSC has emerged as a dominant framework for carrying out species-tree analyses as it naturally accommodates coalescent stochasticity.

Many implementations of the MSC assume that their input is structured into nonrecombining loci or coalescence genes (Ogilvie, Bouckaert and Drummond, 2017; Rannala and Yang, 2003). These “c-genes” (Doyle, 1997; Springer and Gatesy, 2016) correspond to the segments between recombination events in a genome, and therefore each site within a c-gene will share the same phylogenetic history as every other site in the same c-gene. However, in practice these implementations are instead used to analyze data sets structured into m-genes, “a particular sequence of nucleotides along a molecule of DNA…which represents a functional unit of inheritance” (Rieger, Michaelis and Green, 2012; Doyle, 2021). Because this is common practice, we seek to assess the impact of recombination within m-genes on the inference of the divergence times and population sizes by these methods when m-genes are used as input. We will refer to these implementations using “multilocus methods” as shorthand.

There are a few studies to date which have investigated the impact of recombination on species phylogeny estimation. Lanier and Knowles (2012); Wang and Liu (2016) assessed the performance of coalescent-aware methods in the presence of recombination and have shown that species tree topology inference under the MSC appears to be robust to intra-locus recombination. In examining the impact of violations of free inter-locus recombination, Wang and Liu (2016) additionally found that the use of recombination breakpoints for identifying loci improves the accuracy of species tree topology estimation. However, little is known about the extent to which recombination affects the estimation of ancestral population sizes and divergence times. Very recently, Zhu, Flouri and Yang (2022) tested the effects of intralocus recombination on several inference problems including the estimation of population parameters under the MSC. They found that in the “realistic” simulations, Bayesian methods using multilocus sequence data under the MSC performed reasonably well when the amount of recombination was low or moderate, but could underestimate population sizes and inflated species divergence time given elevated recombination rates, confirming similar observations from previous studies (Wall, 2003; Lohse and Frantz, 2014).

The estimation of divergence times along with population sizes of species phylogenies can be conducted in a variety of ways and we choose three modern and representative approaches to study. First, the multilocus method StarBEAST2 (Ogilvie et al., 2017) jointly infers gene and species histories and is an extensively used approach to infer evolutionary parameters while accounting for rate variation and gene discordance. Second, the single nucleotide polymorphism (SNP) method SNAPP (Bryant, Bouckaert, Felsenstein, Rosenberg and RoyChoudhury, 2012) avoids the problem of c-gene/m-gene conflation by assuming the input data set consists of unlinked biallelic markers such as SNPs, since recombination cannot occur within a single site. Third, advances in whole-genome sequencing has been driving the development of another class of methods that use sequentially Markovian approximations of the coalescent to integrate over gene histories and recombination breakpoints (McVean and Cardin, 2005; Steinrücken, Kamm, Spence and Song, 2019; Liu, Ogilvie and Nakhleh, 2021), and we used diCal2 (Steinrücken et al., 2019) to represent this class.

While some degree of model violation is unavoidable for any statistical method when applied to real data, we show that adding recombination and linkage has a similar and mild affect on StarBEAST2 and SNAPP. However, parameters inferred by diCal2 could be wildly erroneous, casting doubt on the approximations employed by that method.

## 2 Material and methods

### 2.1 Simulations

To examine the impact of recombination on continuous parameter estimation given a fixed species topology, we performed whole-genome simulations under a classical coalescent with recombination model using msprime version 1.0.2 (Kelleher, Etheridge and McVean, 2016). We varied the species divergence times, mutation rates, and recombination rates to create data sets approximating different evolutionary scenarios. Specifically, two model species trees with shallow and deep evolutionary timescales were used: i) a shallow phylogeny of height 1.5 million years, and ii) a deep phylogeny of height 18 million years (Fig.1). We considered two per-generation mutation rates, *µ* = 10*−*8 and *µ* = 10*−*7, where the smaller rate is very similar to the rate of 1.1*×* 10^*−*8^ reported by Roach, Glusman, Smit, Huff, Hubley, Shannon, Rowen, Pant, Goodman, Bamshad, Shendure, Drmanac, Jorde, Hood and Galas (2010). We used three different recombination rates: *r* = 2 *×* 10^*−*9^, *r* = 2 *×* 10^*−*8^, and *r* = 2 *×* 10^*−*7^ per site per generation, where the medium rate is similar to the genome-wide average recombination rate of 1.26 cM/Mb reported in Jensen-Seaman, Furey, Payseur, Lu, Roskin, Chen, Thomas, Haussler and Jacob (2004). For each combination of *µ* and *r*, 10 replicates of ten 1-Mb long chromosomal segments were simulated for each taxon. In all cases, the population size was 20,000 diploid individuals, and the generation time was 25 years as in Li and Durbin (2011). DNA sequences were generated assuming the Jukes-Cantor nucleotide substitution model.

### 2.2 Continuous-parameter estimation

For the inference of divergence times and population sizes, the true species tree topology (((A,B),C),D) (Fig. 1) was provided to all methods evaluated here.

**Figure 1:**
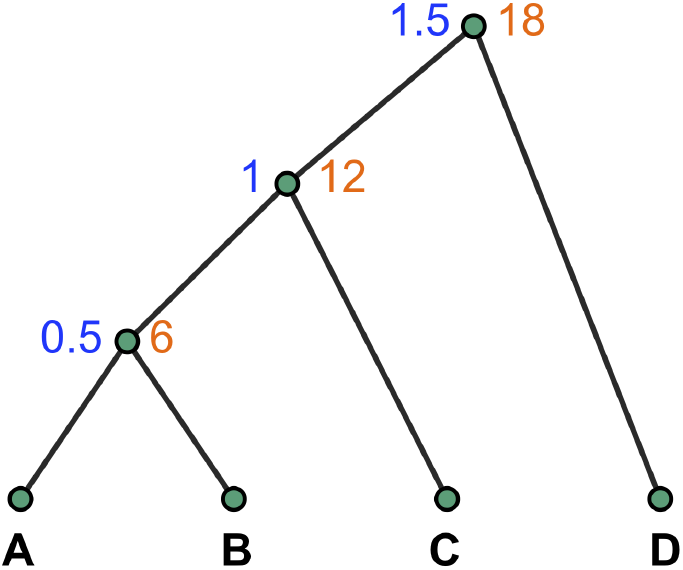
The model species tree used in the simulations. Values in blue and orange at internal nodes represent the node times in million years for the shallow and deep phylogenies, respectively. Population size of 20,000 individuals and generation time of 25 years were assumed.

The Markov chain Monte Carlo (MCMC) convergence of StarBEAST2 and SNAPP was examined by the Gelman-Rubin convergence diagnostic 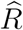 (Gelman and Rubin, 1992), and the effective sample size (ESS).

#### 2.2.1 StarBEAST2

To prepare the sequence alignments for StarBEAST2, five alignments of length 200, 1000, and 5000 bp were extracted from each chromosomal segment by evenly sampling across the segments using python script from https://github.com/ForBioPhylogenomics/tutorials/blob/main/week2_src/make_alignments_from_vcf.py.

The StarBEAST2 analyses used the Jukes-Cantor substitution model with a strict clock where the clock rate was estimated with a 1*/X* prior, and assumed a Yule prior on the species tree. In addition, a normally distributed calibration prior was applied on the root age, with *Normal*(*µ* = 1.5, *σ* = 0.1) for the shallow phylogeny, and *Normal*(*µ* = 18, *σ* = 0.1) for the deep one. Constant population sizes were estimated given a 1*/X* hyperprior (popMean).

The chain length was set to 0.5 billion generations, sampling every 200,000 states, and the first 20% collected samples were discarded as burn-in. Fifteen independent runs (using different seeds) of such chains were carried out on every data set. We generated maximum clade credibility (MCC) trees with posterior mean heights using TreeAnnotator v2.6.6 (Drummond, Suchard, Xie and Rambaut, 2012).

#### 2.2.2 SNAPP

For each data set, the segments from 10 chromosomes were concatenated into one file via BCFtools’s `concat` command (Danecek, Bonfield, Liddle, Marshall, Ohan, Pollard, Whitwham, Keane, McCarthy, Davies and Li, 2021), The SNAPP input XML files were produced by the ruby script snapp prep.rb (Stange, Sánchez-Villagra, Salzburger and Matschiner, 2018). Specifically, the ruby script extracted at most 1000 SNPs, where the minimum distance between SNPs was 1000 nucleotides, with only variable sites included. The root ages were constrained by the same calibration prior as we utilized for the Star-BEAST2 analyses. We used a gamma prior on lambda, whose mean was the true number of speciation events for every substitution per site of tree length. Specifically, the alpha shape parameter was set to 2, and for the shallow phylogney, beta was set to 2500 and 250 on data with low and high mutation rates, respectively. For the deep phylogeny with low and high mutation rates, beta was set to 208.335 and 20.835, respectively. We placed a gamma-distributed prior with an alpha of 8 on theta, whose mean equals 8 *×* 10^*−*4^ and 8 *×* 10^*−*3^ for the low- and high-mutation-rate scenarios, respectively.

MCMC chains were run at least 1 million generations, with the same sampling frequency and burn-in percentage as used in the StarBEAST2 analyses. The chain would be resumed if the ESS of either posterior or likelihood was insufficient (*<* 300) after a completed run. The post-burn-in samples were summarized as an MCC tree with nodes scaled to the mean height estimates using TreeAnnotator v2.6.6 (Drummond et al., 2012).

#### 2.2.3 diCal2

For each generation, we used 70 particles, each of which performed 6 EM-steps, and 6 best points were selected as the parents for the next generation. Time was discretized into intervals by 11 break points, which was chosen log-uniformly between 1000 years and 300 million years. The genetic algorithm was repeated for 5 generations as used in Steinrücken et al. (2019). The parameters with highest composite likelihoods were reported. For fair comparison with other methods with root calibrations, for each replicate, we scaled the parameter estimates of diCal2 such that the root height equals the true value.

## 3 Results

### 3.1 Characteristics of the simulation data

Because the accumulation of recombination is a result of the product of interaction between recombination rate and time span, deep divergence tends to yield shorter c-genes compared with the recent divergence, with a mean length slightly over 62% of that of the shallow phylogeny (Table 1). At both tree depths, recombination rate increases result in shorter c-genes (Table 1). Based on our simulations, we observed that an order of magnitude increase in the recombination rate leads to roughly an order of magnitude decrease in the average c-gene size. As expected, the mutation rate has negligible impact on the lengths of c-genes (results are now shown in the table).

**Table 1:**
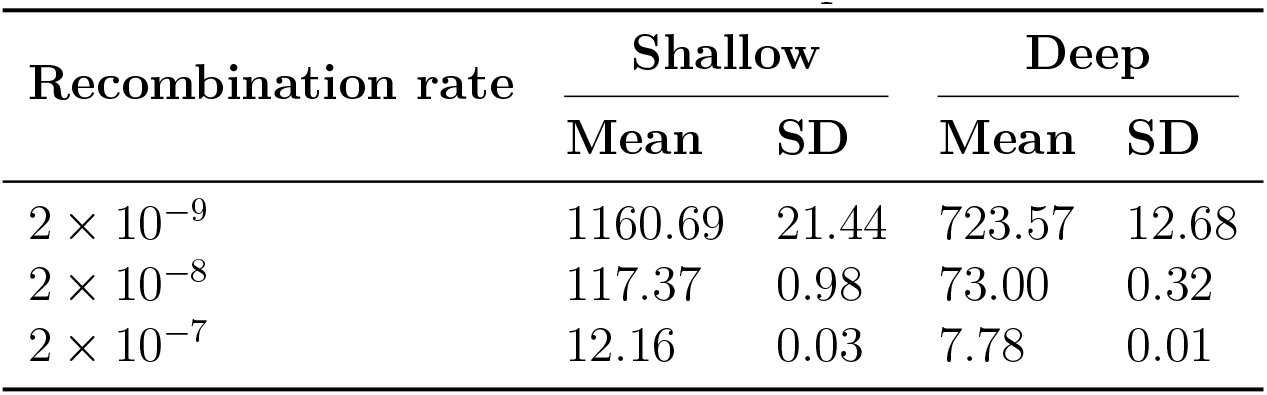
Means and standard deviations of the average c-gene lengths of the simulated data. Each value was obtained from 10 replicates.

| Recombination rate | Shallow |  | Deep |  |
| --- | --- | --- | --- | --- |
|  | Mean | SD | Mean | SD |
| $2 \times 10^{-9}$ | 1160.69 | 21.44 | 723.57 | 12.68 |
| $2 \times 10^{-8}$ | 117.37 | 0.98 | 73.00 | 0.32 |
| $2 \times 10^{-7}$ | 12.16 | 0.03 | 7.78 | 0.01 |

### 3.2 Accuracy of continuous parameter estimates

Taking advantage of the root calibrations, overall, all methods inferred better estimates of divergence times compared with population sizes. Bayesian inference based on the MSC using either linked sites (i.e., StarBEAST2) or SNPs (i.e., SNAPP) performed well for small recombination rates (Supplementary Fig. S1 and Fig. 2).

**Figure 2:**
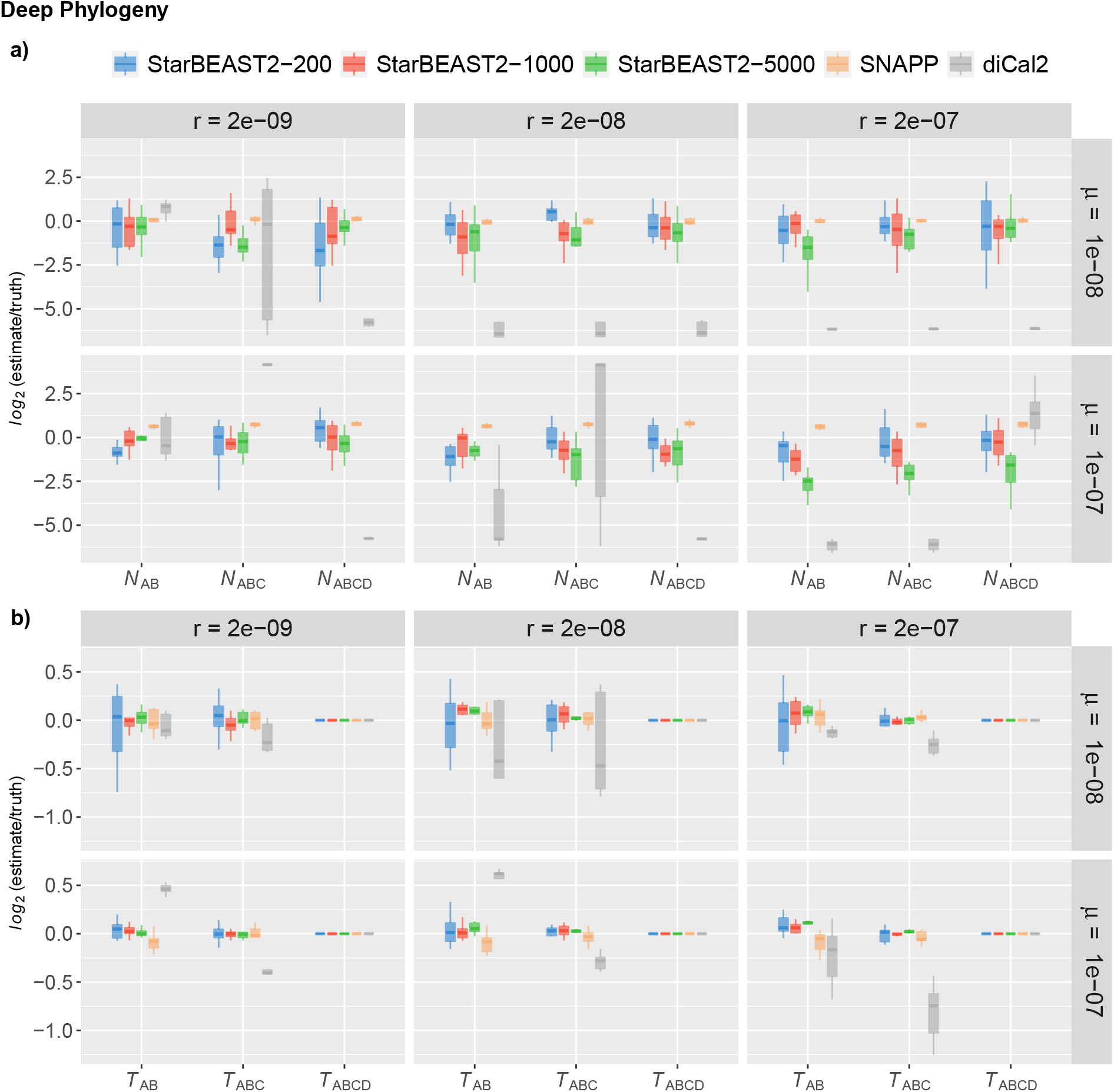
Inference results on the deep phylogeny. Results are shown for five methods: StarBEAST2 with gene alignments of length 200 (StarBEAST2-200), 1000 (StarBEAST2-1000), and 5000 (StarBEAST2-5000), SNAPP, and diCal2. a) Boxplots showing the estimation error of population sizes. *N*_*AB*_ = the population size of the A-B ancestral population, *N*_*ABC*_ = the population size of the A-B-C ancestral population, and *N*_*ABCD*_ = the population size of the root population. b) Boxplots showing the estimation error of divergence times. *T*_*AB*_ = the time of the A-B split, *T*_*ABC*_ = the time of the A-B-C split, and *T*_*ABCD*_ = the time of the A-B-C-D split.

As opposed to StarBEAST2, SNAPP yielded more accurate ancestral population sizes at shallow divergences, reflecting the discussion of Bryant et al. (2012) that “*θ* values can only be reliably inferred for ancestral populations if sufficiently many coalescent events occur within these populations.” We found that, in general, SNAPP had more similar performance to StarBEAST2 supplied with gene of length 200, but with smaller variation. In most cases, larger mutation rates led to lower variance for the results of StarBEAST2 and SNAPP.

It has been previously shown that recombination results in biased estimates of ancestral population parameters (Wall, 2003; Lohse and Frantz, 2014; Zhu et al., 2022). Here we repeated the StarBEAST2 analyses on data with growing segment lengths (200, 1000, and 5000 sites), as longer segments tend to encompass more intra-genic recombination break-points (Table 1). For small recombination rates, more sites slightly improved the estimates of StarBEAST2 (Supplementary Fig. S1 and Fig. 2). As recombination rate increases, the parameter estimates of longer segments started to deviate from the true values, with ancestral population sizes being underestimated but divergence times being overestimated. Remarkably, using sequences with 5000 sites under high recombination rates, StarBEAST2 significantly underestimated the population sizes by 74.5% and 54.6% on average for the shallow and deep species trees, respectively. We noticed that such underestimation of population sizes was accumulated backwards in time for the shallow phylogeny (as reflected by *truth > N*_*AB*_ *> N*_*ABC*_ *> N*_*ABCD*_) but dissipated for the deep phylogeny (as reflected by *N*_*AB*_ *< N*_*ABC*_ *< N*_*ABCD*_ *< truth*). Despite a few exceptions (e.g., *r* = 2 *×* 10^*−*8^, *µ* = 10^*−*8^, and length of 1000 sites), StarBEAST2 using short and intermediate segments seemed to be relatively robust to the presence of recombination. This supports the previous finding that excessive recombination events affected Bayesian analysis of multilocus sequence data assuming the MSC model (Zhu et al., 2022). In addition, the accuracy of StarBEAST2 was slightly higher for data sets that evolved with high mutation rate (*µ* = 10^*−*7^). Such improvement is likely a consequence of more phylogenetic signal contained in the fast-evolving nucleotide data. Furthermore, the estimated population parameters were more accurate and with smaller variance at the deeper timescale.

Although diCal2 was designed to explicitly account for the recombination process, it appears to be the worst-performing one among all the methods evaluated here in this study. We found that diCal2 underestimated the root population size by about 25.9% on average. Notably, we observed a large variance of diCal2’s estimates between replicates for the deep phylogeny when the relative rates of recombination to mutation are 0.2 (*N*_*ABC*_) and 2 (*T*_*AB*_ and *T*_*ABC*_). To determine whether the observed large variance was due to convergence issues, we additionally conducted diCal2 analyses on the full data sets using narrow bounds for the population parameters. We only detected minor convergence problems under a model condition with high recombination rate and mutation rate (*r* = 2 *×* 10^*−*7^, *µ* = 10^*−*7^, and deep phylogeny; Supplementary Fig. S2). Hence, the observed erroneous behavior of diCal2 was not directly tied to convergence issues.

## 4 Discussion

The results presented here show that realistic levels of recombination are likely to have very little impact on Bayesian inference of evolutionary parameters under MSC using either multilocus data or unlinked SNPs. Surprisingly, anomalous behavior of diCal2, which was designed for analyzing full genome sequences in the presence of recombination, was found in our simulation study, most likely caused by extensive approximations employed in its algorithm. Still, the utility of MSC methods will depend on factors including the level of recombination and the lengths of loci. The worse performance of StarBEAST2 when loci had thousands of sites clearly suggests that the amassing of intra-locus recombination events substantially hinders the power of MSC-based multilocus analysis. While diCal2 failed to achieve desirable performance in our study, it is still worth exploring other inference techniques under the multispecies coalescent with recombination that utilize different mathematical or algorithmic frameworks.

Our analyses of how recombination affects the estimation of ancestral population parameters is similar to the recent work by Zhu et al. (2022). Nevertheless, the latter focused on the effect of recombination on Bayesian analyses for addressing different phylogenetic questions under the MSC. In contrast, our work explores the power of different types of methods under various evolutionary scenarios. Zhu et al. (2022) simulated data with intralocus recombination under the multispecies network coalescent (Yu, Dong, Liu and Nakhleh, 2014) model and varied the number of loci as well as number of sequences. We simulated whole-genome data under the MSC, and then sampled distantly-spaced segments of small, medium, and large sizes across the genome, which is a common practice in empirical phylogenomic analyses. As a result, our data included the effect of both intra-locus and inter-locus recombination whereas Zhu et al. (2022)’s data incorporated intra-locus recombination and introgression, but not inter-locus recombination.

It is important to note that neither our study nor Zhu et al. (2022)’s incorporated variation of substitution rates and recombination rates among lineages and sites. Rate heterogeneity has been widely observed in real data (Aguileta, Bielawski and Yang, 2006; Nabholz, Glémin and Galtier, 2008; Auton, Myers and McVean, 2014; Kawakami, Smeds, Backström, Husby, Qvarnström, Mugal, Olason and Ellegren, 2014; Beeson, Mickelson and McCue, 2019). It is definitely worth investigating how reliable current methods are in the presence of varying levels of rate heterogeneity.

Finally, while we limited our study to the impact of recombination on MSC-based species tree inference methods, other evolutionary processes such as gene flow and gene duplication and loss could have an even much larger impact on the performance of those inference methods. In particular, it is worth exploring how the interaction among these processes impact phylogenomic inference under the MSC.

## 5 Funding

This work was supported in part by NSF grants CCF-1514177, CCF-1800723 and DBI-2030604 to L.N.

## Declaration of Competing Interest

The authors declare that they have no known competing financial interests or personal relationships that could have appeared to influence the work reported in this paper.

## Appendix A. Supplementary material

Supplementary data to this article can be found online at

## Notes

### Competing Interest Statement

The authors have declared no competing interest.

